# Safety concerns following the use of ketamine as a potential antidepressant for adolescent rats of both sexes

**DOI:** 10.1101/2024.11.08.622617

**Authors:** Jordi Jornet-Plaza, Sandra Ledesma-Corvi, M. Julia García-Fuster

## Abstract

While ketamine is already approved for treatment resistant depression in adult patients, its efficacy and safety profile for its use in adolescence still needs further investigations. Preclinical studies proved dose- and sex-dependent effects induced by ketamine during adolescence, but few studies have evaluated the short- and long-term safety profile of ketamine at the doses necessary to induce its antidepressant-like effects. The present study aimed at evaluating the antidepressant-like effects of ketamine (1, 5 or 10 mg/kg; vs. vehicle; 1 vs. 7 days) during adolescence in naïve or early-life stressed (i.e., maternal deprivation) rats of both sexes in the forced-swim or novelty-suppressed feeding tests. Safety was evaluated by measuring the psychomotor- and reinforcing-like responses induced by adolescent ketamine. In addition, long-term safety was evaluated in adulthood at the level of cognitive performance, or addiction liability (induced by a challenge dose of ketamine in rats treated with adolescent ketamine). The main results reinforced the potential for ketamine as an antidepressant for adolescence, but at different dose ranges for each sex. However, some safety concerns emerged for adolescent female rats (i.e., signs of sensitization at the dose used as antidepressant) and adult male rats (i.e., addiction liability when re-exposed to ketamine in adulthood), suggesting the need for caution and further research before moving forward the use of ketamine as an antidepressant for adolescence.

## Introduction

Major depression is not a disorder exclusive to adulthood, since it also affects the adolescent population, being the most common affective-like disorder with a prevalence of 5-6% in teenagers (e.g., [1]) that is rising in the last years (reviewed by [2]). Unfortunately, the therapeutic options for adolescent depression are limited, with fluoxetine or escitalopram as the recommended choices in combination with psychological therapy [3-5]. This scarce number of safe pharmacological options might be related to the observed age differences in the pathophysiology of the disorder (e.g., [6]), since the adolescent brain is still under development, with antidepressants showing a lower response as the one induced in adulthood [7]. In addition, given that adolescent depression is characterized by an elevated risk of suicidal behaviors (e.g., [8-9]), there is an urgent need to characterize novel fast-acting antidepressants for this vulnerable age group (recently reviewed by [10]).

From this perspective, ketamine, an NMDA receptor antagonist, was approved by the FDA in 2019, followed by several other European countries, as a fast-therapeutic approach in adult patients with treatment resistant depression (reviewed by [11]). The use of ketamine as an antidepressant is recommended, in combination with a classical antidepressant, only in severe cases of resistant depression in adults. Therefore, ketamine seems like a good candidate to be further explored as a fast-acting antidepressant for adolescence. In fact, numerous clinical trials have already showed favorable results in terms of its antidepressant efficacy in adolescence [12-15] (reviewed by [16]). However, there are still a lot of unknowns regarding the potential adverse effects of ketamine on the developing brain and the long-term consequences of its use in adolescence, so safety evaluations are urgently needed (as discussed by [17]).

In this sense, one of the negative effects that might be associated with ketamine’s administration during adolescence is its abuse potential (e.g., [18-19]). The fact that ketamine is a recreational drug (e.g., [20-21]) and that an early age of drug exposure is a variable that has a major influence on future consumption (e.g., [22]) makes it a major topic of discussion. Some examples include several preclinical studies that showed that ketamine administered at subanesthetic doses was able to induce conditioning in mice [23-24]. Also, a more recent study revealed some sex and dose dependent effects of ketamine’s reinforcing properties at doses used for treating various psychopathologies, as well as the capability of ketamine to induce similar reinstatement-rates in both sexes [25]. Yet, previous studies regarding the additive-like potential of ketamine at subanesthetic doses are scarce, especially during adolescence, since most of the studies found in the literature centered in adult rodents.

Although recent clinical studies described that ketamine’s administration at subanesthetic doses for treatment-resistant depression in adulthood is safe in terms of cognition [26-27], some concerns have been described for ketamine when used as a recreational drug (i.e., deficits in cognition and working memory; see [28-30]). Moreover, most of these studies were conducted in adulthood, so little information is available regarding the impact of adolescent ketamine on cognitive performance. In fact, the few studies that are available presented variable results. Either ketamine showed no long-term effects on cognition or reward processing (e.g., [31]), or displayed long-term cognitive deficits after repeated treatment during pre-adolescence [32].

Against this background, the present study aimed at evaluating potential safety concerns associated with the use of ketamine as an antidepressant in adolescence while incorporating sex as a biological variable. To do so, the antidepressant-like efficacy of different doses of ketamine was evaluated in adolescent rats, as well as its reinforcing- and psychomotor-like responses. Moreover, long-term safety in adulthood following the adolescent treatment was evaluated at the level of cognitive performance and addictive-like potential (i.e., rewarding-like responses following an acute ketamine challenge in adulthood).

### Experimental procedures

#### Animals

All experimental procedures were approved by Local Bioethical Committees (CEEA 155-12-20 and 2021/’05/AEXP; Conselleria Medi Ambient, Agricultura i Pesca, Direcció General Agricultura i Ramaderia, Govern de les Illes Balears), in accordance to the ARRIVE guidelines [33] and following the EU Directive 2010/63/EU. A total of 332 Sprague-Dawley rats (167 males, 165 females) were utilized in different experimental procedures as detailed in Fig. 1. Rats were bred in the animal facility at the University of the Balearic Islands. While some of these rats were used in naïve conditions (102 males and 100 females; Fig. 1A-C) others were exposed to early-life stress (65 males and 65 females; Fig. 1D). In particular, whole litters were subjected to maternal deprivation in their home cage from post-natal day (PND) 9 to PND 10 (a single 24 h episode) with no nutritional supplements, while their mothers stayed in nearby cages as previously executed [34]. All pups were weighted before and after the procedure.

**Fig. 1.**
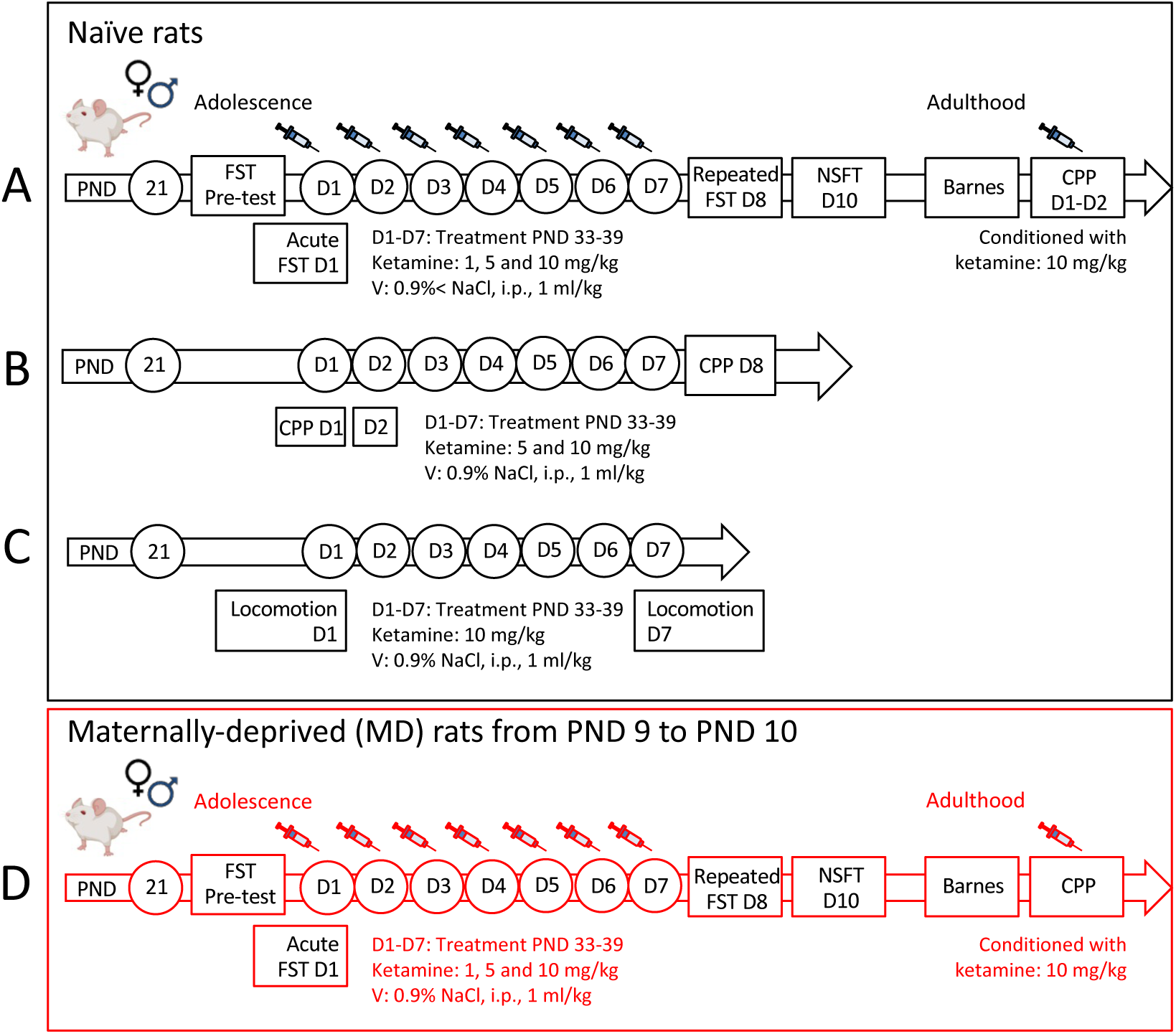
Experimental timeline. Effects induced by different doses of ketamine (1, 5 or 10 mg/kg) in **(A-C)** naïve and **(D)** maternally deprived (MD) adolescent rats of both sexes. Experimental procedures to evaluate potential antidepressant-like effects of ketamine during adolescence and its long-term safety in adulthood (i.e., cognitive performance and reinforcing-related responses) in **(A-C)** naïve and **(D)** MD rats of both sexes. Experimental procedures to evaluate potential reinforcing-related responses of ketamine (5 or 10 mg/kg) **(B)** as well as possible psychomotor effects to the highest dose of ketamine tested (10 mg/kg) **(C)** in adolescent naïve rats of both sexes. CPP: conditioned place preference; D: day; FST: Forced swim test; NSFT: novelty suppressed feeding test; PND: post-natal day; V: vehicle.

Rats from all studies were separated at weaning and housed in standard cages (2-4 rats/cage/sex) with unlimited access to a standard diet and water in a controlled environment (22 °C, 70% of humidity, and a 12:12 h light/dark cycle, lights on at 8:00 AM). All procedures were performed during the light-period and all efforts were directed towards minimizing the number of rats used, the number of procedures and their suffering. In line with our prior studies (e.g., [35-36]) and to avoid unnecessary stress in females, the particular phases of the estrous cycle were not examined, since cyclicity was not part of our research query [37] and both sexes seemed equally variable due to hormonal periodicity [38-39]. In fact, the observed individual variability for each behavioral measure analyzed in this study for males and females reinforced that notion.

#### Ketamine treatment during adolescence

As detailed in Fig. 1, rats were treated throughout a period during mid and late-adolescence [40] for 7 consecutive days (1 injection per day, i.p., 1 ml/kg, from PND 33-39, and as previously done, see [35]) with ketamine (Anesketin: 100 mg/ml of ketamine from Dechra Pharmaceuticals, Northwich, United Kingdom; doses: 1, 5 and 10 mg/kg) or vehicle (0.9% NaCl). The dose-range was selected from previous studies form our group [35-36] and others (e.g., [41]).

#### Behavioral screening during adolescence

##### Antidepressant-like responses of adolescent ketamine

Antidepressant-like responses were ascertained by diverse tests previously validated in the field (Fig. 1A and 1D). We first screened the antidepressant-like response induced by ketamine under the stress of the forced-swim test in adolescent naïve and maternally-deprived rats of both sexes, since this test has been the goal standard screening tool in the industry for antidepressant-like responses. Following standard procedures [42], slightly modified in our group (e.g., [43-44]), rats were exposed to a 15-min pre-test session in which they were individually placed in water tanks (41 cm high x 32 cm diameter, 25 cm depth; temperature of 25 ± 1 °C), so they could learn that no escape was available. The typical behavioral responses compare immobility vs. activity (climbing or swimming) times. The next day, 30 min post-treatment (Fig. 1A and 1D), rats were exposed to the water tanks for the actual test that lasted 5-min and during which rats were videotaped. Moreover, given that similar repetitive screening testing provided prior reliable measurements across time (see [45,35]), rats were individually re-scored in this test 24 h after the last repeated treatment dose (on D8; see Fig. 1) for a 5 min session that was also videotaped. Videos were later evaluated by an experimenter blind to the particular treatment conditions with Behavioral Tracker software (CA, USA). The time each rat spent (s) immobile was used as an indicative of behavioral despair, while the active time (swimming or climbing) suggested escaping-like behaviors and are indicatives of an antidepressant-like response. To avoid potential behavioral interferences caused by individual excrement samples, water was changed for each animal.

To complement the results from the forced-swim test, rats were also scored in the novelty-suppressed feeding test, which captures antidepressant-like responses under a stressful situation (e.g., [46]). In particular, rats were food-deprived for 48 h, since motivation for food is required for this particular test, and then individually placed in a square open arena (60 cm x 60 cm, and 40 cm in high) under housing lighting conditions with three food pellets in the center and allowed to freely explore during 5 min [34-35]. The test was performed 3 days post-treatment (Fig. 1A and 1D). Sessions were videotaped to then analyze feeding time (s), and latency to center (s). To avoid potential behavioral interferences caused by individual odors the arena was cleaned with 70% ethanol in between animals.

##### Rewarding-like responses of adolescent ketamine

The conditioning protocol of a single dose of ketamine that we followed was based on the design previously described by [47] lasting 2 consecutive days. Each day, rats were moved to the procedural room and allowed to acclimate for 1 hour. The behavioral test was performed in an apparatus with two visually different chambers (30 x 30 cm), one with the walls with black stripes and the metal floor with square holes, and the other one with the walls with black circles and the metal floor with circular holes. The two chambers were separated by a central corridor (10 x 30 cm) without any visual cues, and connected through sliding doors. On day 1 (CPP D1, PND 33; Fig. 1B), all rats were administered vehicle (0.9% NaCl, 1 ml/kg, i.p.), and placed in one of the randomly assigned compartments where the animal was confined for 20 minutes. After 3 hours, rats were treated with either ketamine (5 or 10 mg/kg, i.p.) or vehicle (depending on the experimental group) and confined in the other compartment for 20 minutes. Chambers were randomly paired with saline or ketamine to avoid a place preference. The next day (CPP D2, PND 34; Fig. 1B), rats were placed in the central area of the apparatus and were allowed to freely explore the 3 compartments (paired-chamber, central zone and unpaired-chamber) for 20 minutes while the session was recorded. After that, the repeated ketamine paradigm was continued with a daily injection (on D2 the injection was right after the test was finished) until D6 of treatment (PND 38). Finally, on PND 39, a dose of saline was administered again followed by 3 hours later, an injection with a dose of saline or ketamine (5 or 10 mg/kg) i.p., thus completing the 7 doses of saline or ketamine (depending again on the treatment group). Rats paired their treatment (saline vs. ketamine) in the same chambers as performed on PND-33. The next day (PND 40; Fig. 1B), rats were placed in the central area of the apparatus and were allowed to freely explore the 3 compartments (paired-chamber, central zone and unpaired-chamber) for 20 minutes while the session was recorded. The time spent by each animal in each compartment was analyzed (SMART Video Tracking Software, Panlab Harvard Apparatus) and the % time spent in the paired chamber, the number of entries in the paired chamber and the distance traveled (cm) was calculated for rats exposed to 1 or 7 doses of ketamine.

##### Psychomotor-like responses of adolescent ketamine

The effects of a single dose (10 mg/kg, i.p., D1) or repeated doses (10 mg/kg, 7 days, 1 dose/day, i.p.) of ketamine (Fig. 1C) were scored in adolescent rats in an open field arena (85 x 54 cm) during 60 min post-injection. Briefly, on day 1 (D1, PND 33; Fig. 1C), animals were allowed to habituate to the open field for 30 min. Next, rats were treated with a single dose of ketamine (10 mg/kg, i.p.) or vehicle (0.9% NaCl, 1 ml/kg, i.p.) and placed in the open field again. Behavior was recorded for 60 minutes. After that, the repeated ketamine treatment was continued with a daily injection until D6 of treatment (PND 38). Finally, on day 7 (D7, PND 39; Fig. 1C) a dose of ketamine (10 mg/kg, i.p.) or vehicle was administered, thus completing the 7 doses of saline or ketamine (depending on the treatment group) and the locomotion test was performed again. The distance traveled (cm) during 60 minutes after 1 or 7 doses of ketamine was measured for each animal using the software (SMART Video Tracking Software, Panlab Harvard Apparatus), as well as the total accumulated distance (cm) over the 60-min that lasted the test.

#### Behavioral screening during adulthood following adolescent ketamine exposure

##### Cognitive-like responses

The effects of the adolescent ketamine treatment (1, 5, 10 mg/kg, 7 days, PND 33-39, Fig. 1A and 1D) on the long-term cognitive performance of rats was evaluated in the Barnes maze. Briefly, the Barnes maze used in the experiment was a circular platform with 18 holes evenly spaced around its perimeter, with one hole leading to an escape box or target below. The room where the test was conducted had visual cues to provide spatial references for rats to locate the escape box. A bright light served as an aversive stimulus to motivate rats to find the target, leveraging their natural agoraphobia. On the first day, rats were habituated to the maze by placing them in a black start chamber located at the center of the maze under a bright light (500 W). After 10 seconds, the chamber was lifted, and rats were allowed 3 minutes to find and enter the black escape box. On the test day, each rat underwent three training trials, each separated by 10 minutes. Each trial ended when the rat entered the target box or after 3 minutes, at which point the rat was manually placed in the target box and left there for 1 minute to habituate. Ten minutes after the training trials, the actual test began, allowing rats to freely explore the maze for 90 seconds to find the target box. This test was repeated 24 hours later. The amount of time spent (s) to resolve the maze, as well the time progression, were used as a measure of spatial working memory performance (e.g., [48,44]).

##### Rewarding-like responses following an acute ketamine challenge in adulthood

The effects of the adolescent ketamine treatment (1, 5, 10 mg/kg, 7 days, PND 33-39, Fig. 1A and 1D) on the long-term rewarding-like effects induced by an acute challenge with ketamine was evaluated in adult rats with the conditioned place preference test. Briefly, and following a similar paradigm as the one described above, the first day in adulthood rats received a dose of vehicle (0.9% NaCl, i.p.) followed by, 3 hours later, a single dose of ketamine (10 mg/kg, i.p.) or vehicle (Fig. 1A and 1D). Chambers were randomly paired with saline or ketamine to avoid a place preference. The next day (Fig. 1A and 1D), rats were placed in the central area of the apparatus and were allowed to freely explore the 3 compartments (paired-chamber, central zone and unpaired-chamber) for 20 minutes while the session was recorded. The time spent by each animal in each compartment was analyzed (SMART Video Tracking Software, Panlab Harvard Apparatus) and the % time spent in the paired chamber, the number of entries in the paired chamber and the distance traveled (cm) was calculated for rats exposed to a repeated paradigm of ketamine in adolescence and challenged with an acute dose of ketamine in adulthood.

### Statistical analysis

Data was analyzed and graphs were plotted with GraphPad Prism, Version 10 (GraphPad Software, CA, USA). Following the guidelines for reporting data and statistical results in experimental pharmacology [49-50], results are displayed as box and whiskers incorporating min to max values and showing symbols for individual values for each rat. Two-way ANOVAs were mainly used for statistical analysis of the data, with Sex and Treatment as the independent variables, except for the locomotor response induced by ketamine which was analyzed across time through three-way ANOVAs (Sex, Treatment, Time) and cognitive performance during the training sessions in the Barnes maze that used Session and Treatment as the independent variables. The particular tests used are detailed in the Supplementary Materials (Tables S1, S2 and S3). Post-hoc comparisons were performed when appropriate. The level of significance was set at *p* ≤ 0.05.

## Results

### Antidepressant-like effects of ketamine in adolescence

Ketamine induced signs of an antidepressant-like response in naïve rats of both sexes, but at different doses and regimens of administration (see Supplementary Table S1 for the particular statistical results). In particular, acute ketamine (dose of 10 mg/kg) induced a significant reduction in immobility as observed 30 min post-ketamine administration in female naïve rats (-57 ± 12 s, \*\*\**p* < 0.001 vs. vehicle-treated rats), which paralleled an increase in climbing behavior (+53 ± 11 s, \*\*\**p* < 0.001 vs. vehicle-treated rats; data not shown). The repeated treatment with ketamine (7 days of a daily dose injection) in adolescent naïve rats, showed signs of antidepressant-like responses, such as the ones observed in the forced-swim test as measured 1-day post-treatment (decreased immobility by the dose of 10 mg/kg: -17 ± 7 s, \**p* = 0.032; Fig. 2B). No significant changes were induced by acute or repeated ketamine in adolescent male naïve rats in the forced-swim test (Fig. 2A-B and Supplementary Table S1). However, the novelty-suppressed feeding test, performed 3 days post-treatment, showed signs of improvements induced by the dose of 5 mg/kg of ketamine in the time spent feeding both for male (+28 ± 10 s, \**p* = 0.011) and female rats (+27 ± 9 s, \**p* = 0.014; Fig. 2C) as compared to vehicle-treated rats (Fig. 2C). No other effects were observed in this test (i.e., latency to center, distance travelled; data not shown). Overall, ketamine induced signs of efficacy for both sexes but at different dose-ranges, in line with observed overall Sex differences observed (see Supplementary Table S1).

**Fig. 2.**
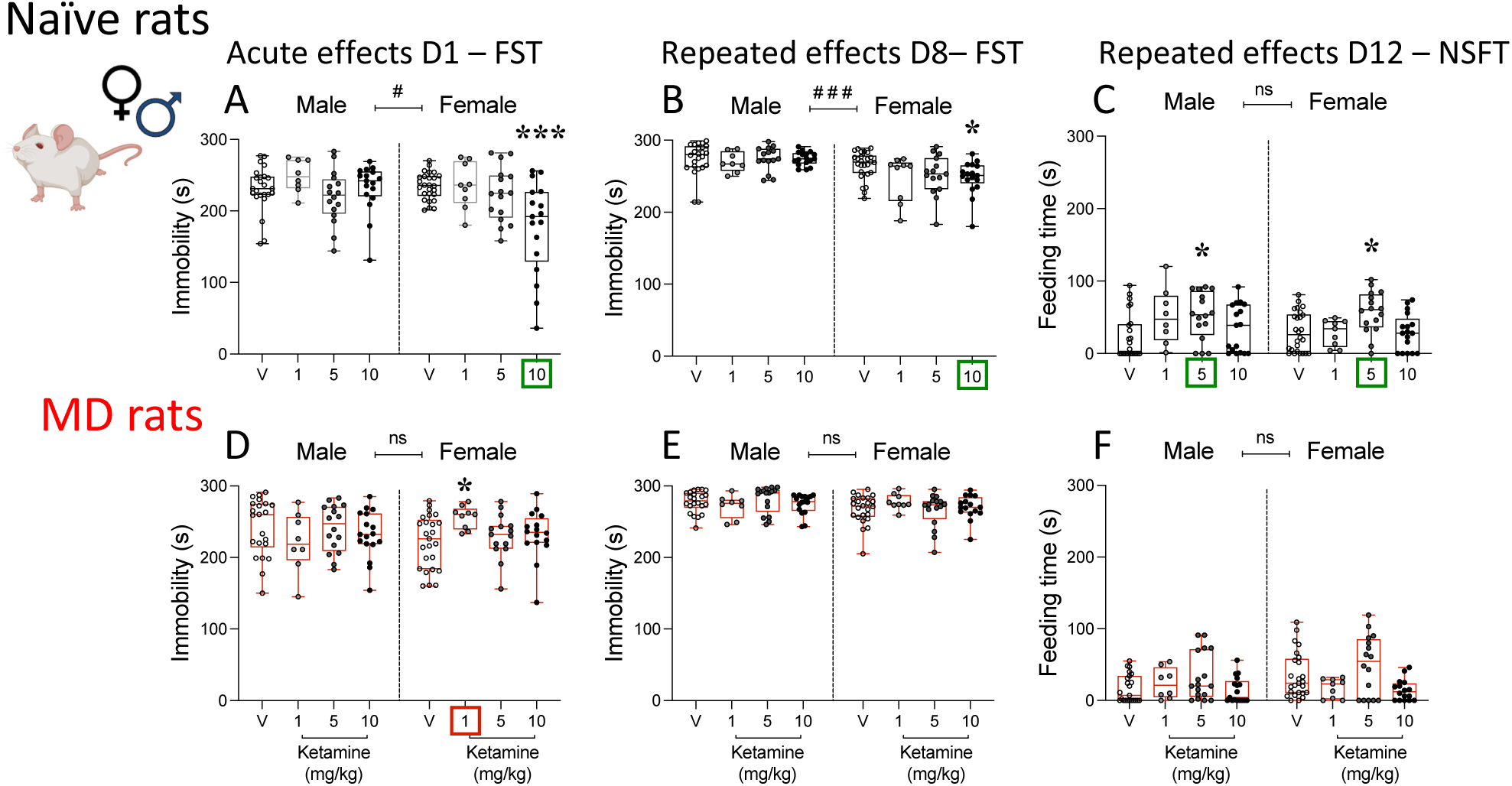
Antidepressant-like effects of ketamine in adolescent rats of both sexes. Acute effects of ketamine (1, 5 and 10 mg/kg, i.p.) as measured 30 min post-treatment on Day 1 (D1) in the forced-swim test (FST) in adolescent **(A)** naïve and **(D)** maternally-deprived (MD) rats. Repeated effects of ketamine (1, 5 and 10 mg/kg, i.p., 7 days, 1 dose/day) as measured 1-day post-treatment on D8 in the FST in adolescent **(B)** naïve and **(E)** MD rats. Repeated effects of ketamine (1, 5 and 10 mg/kg, i.p., 7 days, 1 dose/day) as measured 3-days post-repeated treatment on D10 in the novelty-suppressed feeding test (NSFT) in adolescent **(C)** naïve and **(F)** MD rats. Data represent mean ± SEM of the time spent (s) immobile **(A, B, D, E)** or feeding **(C, F)**. Individual values are shown for each rat. Two-way ANOVAs (independent variables: Sex and Treatment) are shown in Supplementary Table S1. Significant effects of Sex: ###*p* < 0.001 and #*p* < 0.05 when comparing male vs. female rats. \*\*\**p* < 0.001 and \**p* < 0.05 vs. the corresponding Vehicle (V) group.

Interestingly, when ketamine was administered in rats previously exposed to maternal separation early in life, no signs of efficacy were observed for any doses of behavioral tests performed for male or female adolescent rats (see Fig. 2D-E). In fact, 1 mg/kg of ketamine even increased immobility in female rats (+35 ± 11 s, \**p* = 0.031 vs. vehicle-treated rats), showing deleterious signs (Fig. 2D).

### Reinforcing-like effects of ketamine in adolescence

Ketamine (5 and 10 mg/kg) did not induce changes in the conditioned-place preference test, as measured by the % time spent in the paired-chamber, the number of entries in the paired chamber and/or the distance travelled for naïve adolescent rats of both sexes (Fig. 3 and Supplementary Table S1). These lacks of conditioning effects were observed both following an acute (Fig. 3A) or repeated (Fig. 3B) treatment in adolescence.

**Fig. 3.**
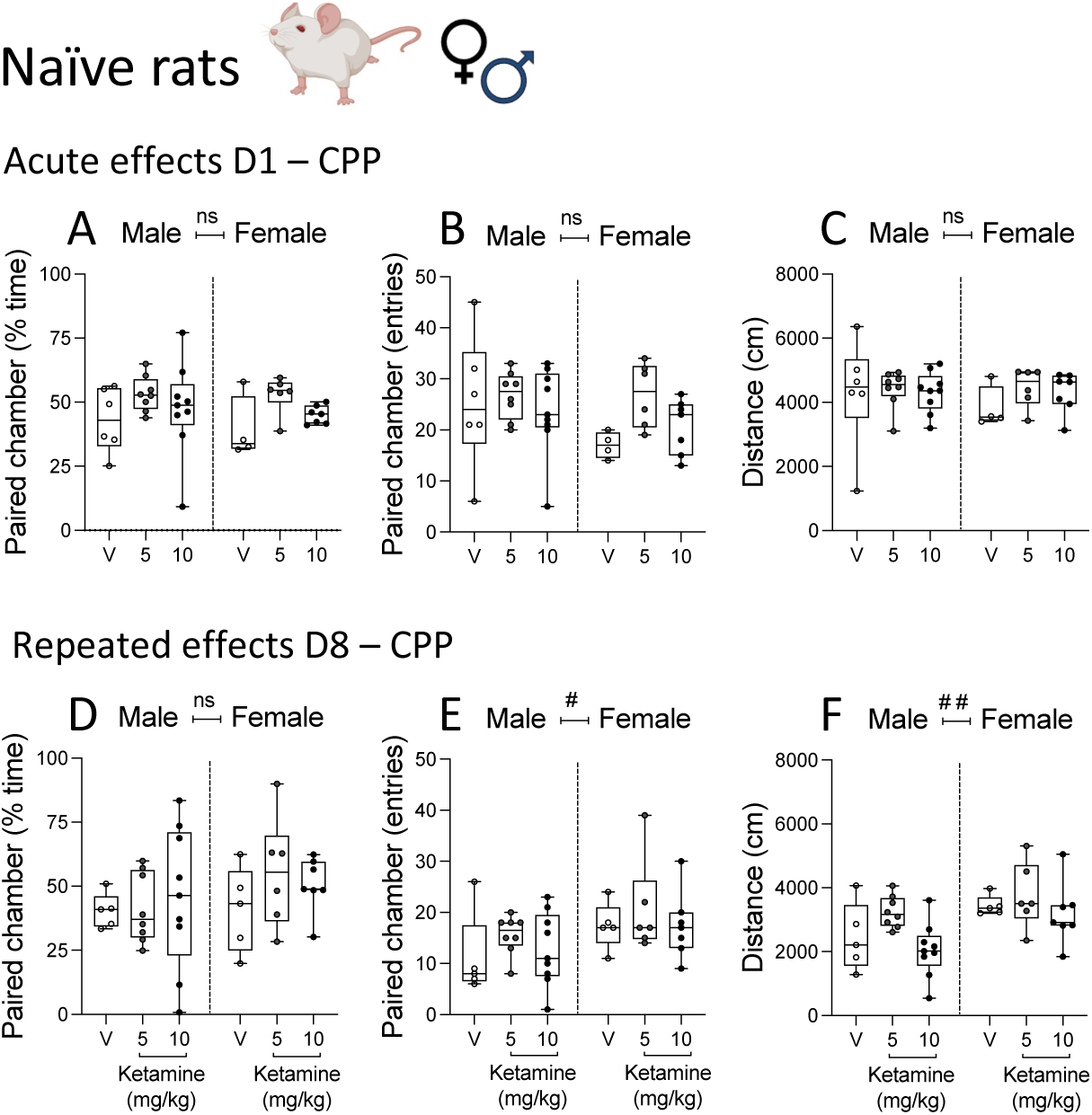
Reinforcing-like effects of ketamine in adolescent rats of both sexes. **(A-C)** Acute effects exerted by a single dose of ketamine (5 and 10 mg/kg, i.p.) exposure in the conditioned-place preference test (CPP) in adolescent naïve rats. **(D-F)** Repeated effects of ketamine (5 and 10 mg/kg, i.p., 7 days, 1 dose/day) in the CPP in adolescent naïve rats. Data represent mean ± SEM of the % time spent in the paired chamber **(A, D)**, the number of entries in the paired chamber **(B, E)**, and the distance (cm) traveled during the test **(C, F)**. Two-way ANOVAs (independent variables: Sex and Treatment) are shown in Supplementary Table S1. Significant effects of Sex: ##*p* < 0.01 and #*p* < 0.05 when comparing male vs. female rats.

### Psychomotor-like effects of ketamine in adolescence

The psychomotor effects of ketamine were evaluated after 1 (D1) and 7 doses (D7), right after treatment and for 1 h in adolescent naïve rats of both sexes. The statistical analysis (see Supplementary Table S2) for both days (D1 and D7) showed significant effects of Sex (i.e., overall higher locomotion for female rats), Treatment (i.e., ketamine increased locomotion) and Time (i.e., increased effects right after treatment). Tukey’s *post-hoc* comparisons revealed that ketamine increased locomotion in a time-dependent manner in female rats after an acute dose (D1: 0-5 min: +949 ± 169 cm, \*\*\**p* < 0.001; 5-10 min: +688 ± 169 cm, \**p* = 0.043; Fig. 4A) or 7 doses (D7: 5-10 min: +2463 ± 371 cm, \*\*\**p* < 0.001; 10-15 min: +1888 ± 371 cm, \*\*\**p* < 0.001; Fig. 4B) vs. vehicle-treated rats. After 15 min the activating effects of ketamine reverted to normal. No significant changes were observed in male rats (Fig. 4A-B).

**Fig. 4.**
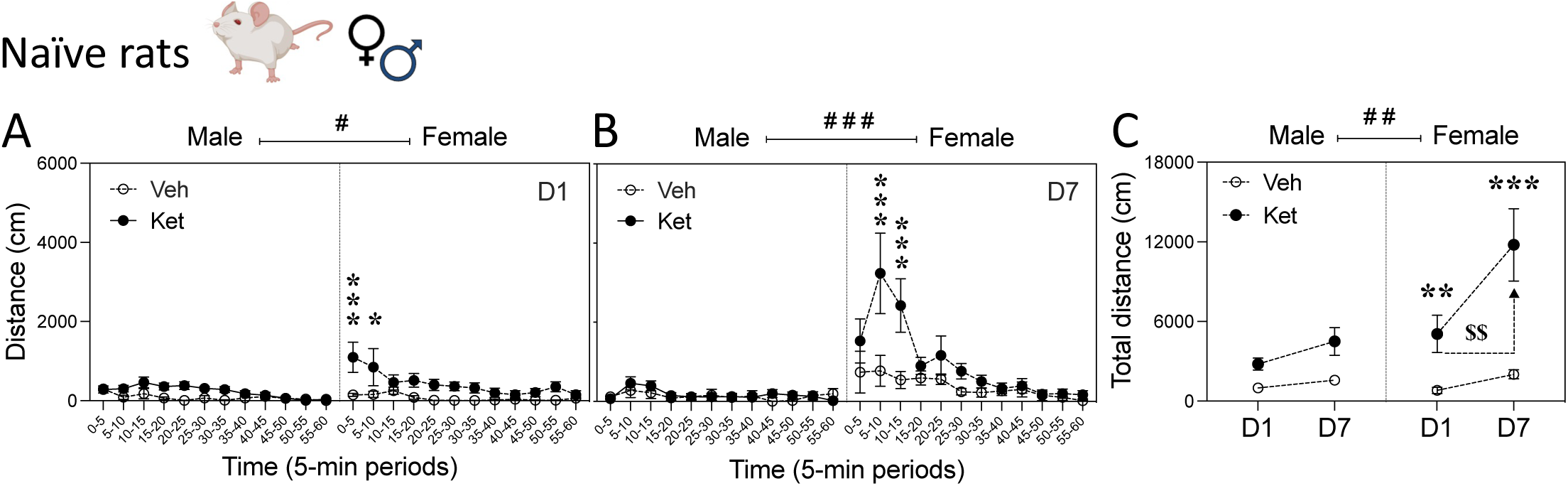
Psychomotor-like effects of ketamine in adolescent rats of both sexes. **(A)** Acute effects exerted by a single dose of ketamine (10 mg/kg, i.p.) exposure in adolescent naïve rats as measured in an open field (Day 1, D1). **(B)** Repeated effects exerted by ketamine (10 mg/kg, i.p., 7 days, 1 dose/day) as measured in an open field right after the last dose in adolescent naïve rats (D7). Data represent mean ± SEM of the distance travelled (cm) in periods of 5-min post injection (locomotion analyzed for 1 h). Three-way ANOVAs (independent variables: Sex, Treatment and Time) are shown in Supplementary Table S2. Significant effects of Sex: ###*p* < 0.001 and #*p* < 0.05 when comparing male vs. female rats. \*\*\**p* < 0.001 and \**p* < 0.05 vs. the corresponding Vehicle (V) group. **(C)** Total distance traveled (cm) when measured after the acute (D1) and repeated (D7) treatments in adolescent naïve rats of both sexes. Data represent mean ± SEM of the accumulated distance travelled (cm) during 60-min post injection. Three-way ANOVAs (independent variables: Sex, Treatment and Day) are shown in Supplementary Table S2. Significant effects of Sex: ##*p* < 0.01 when comparing male vs. female rats. Significant effects of Day: $$*p* < 0.01 when comparing D7 vs. D1. \*\*\**p* < 0.001 and \*\**p* < 0.01 vs. the corresponding Vehicle (Veh) group.

When comparing the total accumulative distance travelled during the 1 h that was monitored, the results showed a significant effect of Sex (i.e., overall higher locomotion for female rats), Treatment (i.e., overall increased effects by ketamine) and Day (higher locomotor responses on D7 than D1; Fig. 4C and Supplementary Table S2). *Post-hoc* analysis confirmed the psychomotor activating effect of ketamine in female rats both at D1 (+4252 ± 1082 cm; \*\**p* = 0.001) and D7 (+9753 ± 2393 cm; \*\*\**p* < 0.001), plus a sensitized response over time as observed when comparing the higher response of D7 (+6709 ± 1672 cm; $$*p* = 0.002) vs. D1 (Fig. 4C).

### Long-term effects of adolescent ketamine on cognitive performance in adulthood

Cognitive performance was evaluated in adult rats in the Barnes maze. The results proved that maternal separation worsened cognitive performance in adulthood as compared to naïve rats, during training and test sessions (see Supplementary Fig. S1). However, ketamine treatment (1, 5 and 10 mg/kg) during adolescence did not induce any long-term changes in cognitive performance neither in naïve (Fig. 5A-C) or maternally-deprived (Fig. 5D-F) rats in adulthood as compared to adolescent vehicle-treated rats (see Supplementary Table S3).

**Fig. 5.**
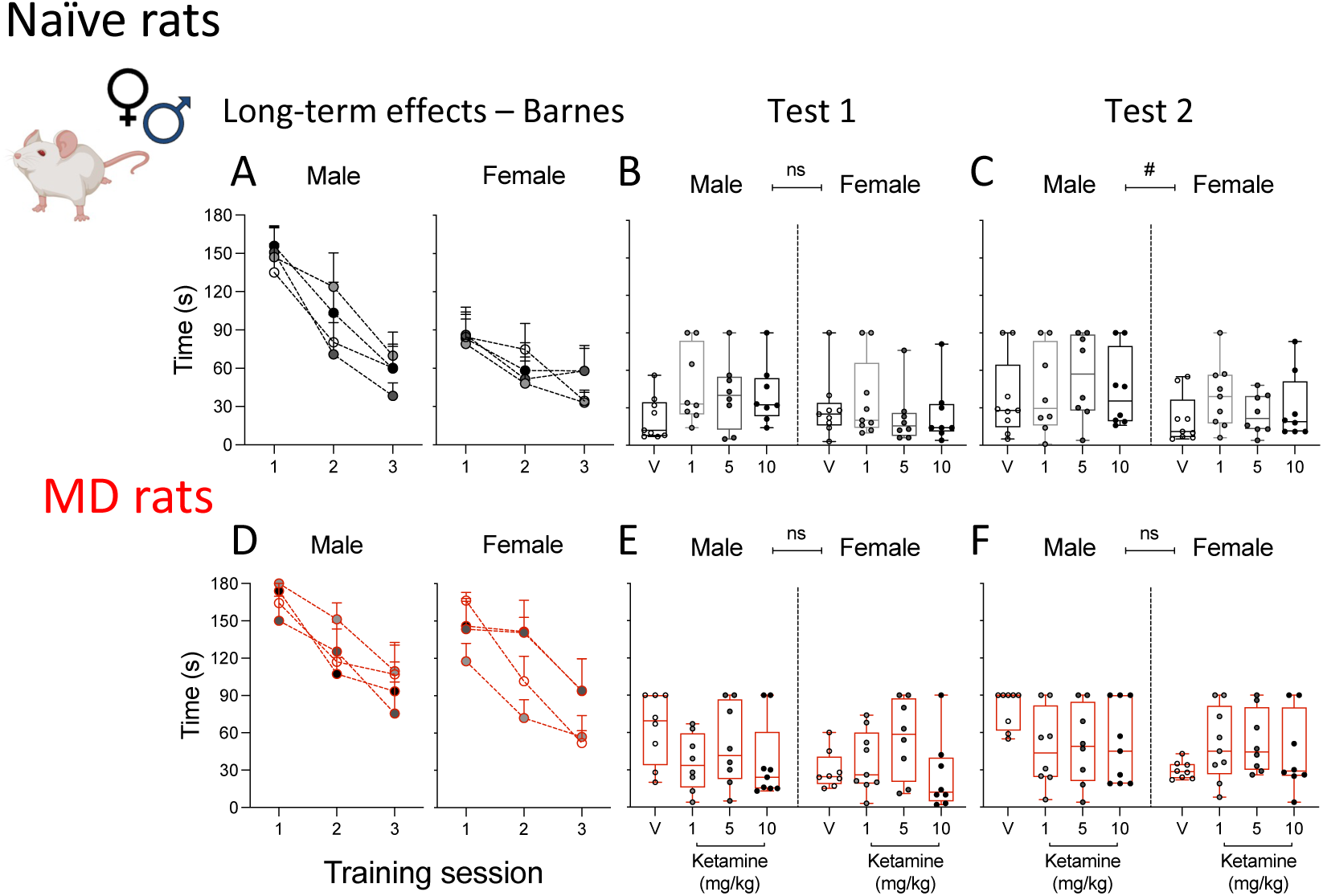
Long-term effects of adolescent ketamine on cognitive performance in adult rats of both sexes. Long-term effects of adolescent ketamine exposure (1, 5 and 10 mg/kg, i.p., 7 days, 1 dose/day) as measured in adult rats in the Barnes maze test in **(A-C)** naïve and **(D-E)** maternally-deprived (MD) rats. Data represent mean ± SEM of the time spent (s) to resolve the test during 3 training sessions **(A, D)**, Test 1 **(B, E)**, and Test 2 **(C, F)**. Individual values are shown for each rat. While training sessions lasted 180 s, test sessions were 90 s each. Test 2 was performed 24 h after the first test with different cues. The progression of the response during the training session was evaluated through two-way ANOVAs (independent variables: Session and Treatment) for each Sex (Supplementary Table S3). Please note that the significant effect of Session (i.e., learning process across sessions) is not shown in graph. Tests’ performances (Test 1 and Test 2) were analyzed through two-way ANOVAs (independent variables: Sex and Treatment; Supplementary Table S3). Significant effect of Sex: #*p* < 0.05 when comparing male vs. female rats.

### Long-term effects of adolescent ketamine on a later challenge with acute ketamine in adulthood

The reinforcing properties of a 10 mg/kg dose of ketamine were evaluated in the conditioned-place preference in adulthood in rats previously exposed to either ketamine (1, 5 or 10 mg/kg) or vehicle in adolescence. In naïve rats, ketamine did not increase the % time spent in the paired chamber (Fig. 6A), nor the number of entries (Fig. 6B), but exerted a significant Sex x Treatment interaction when analyzing the distance travelled (Fig. 6C; see Supplementary Table S3). In fact, *post-hoc* comparisons revealed that adolescent ketamine (1 mg/kg: +940 ± 367 cm; \**p* = 0.036; 5 mg/kg: +942 ± 367 cm; \**p* = 0.035) increased the distance travelled in adult female rats when challenged with a 10 mg/kg dose of ketamine (Fig. 6C).

**Fig. 6.**
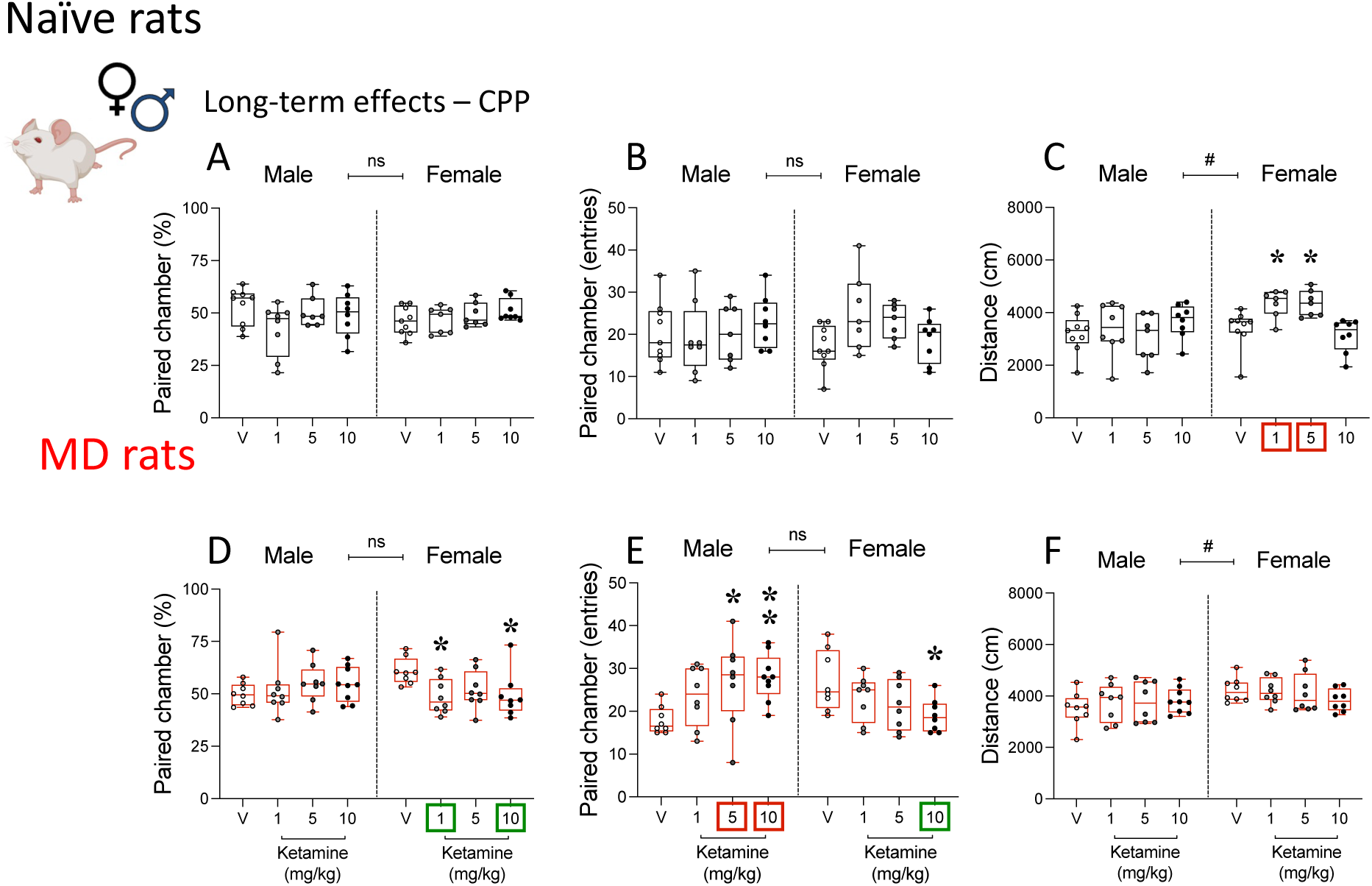
Long-term effects of adolescent ketamine on a later challenge with acute ketamine in adult rats of both sexes. Long-term effects of adolescent ketamine exposure (1, 5 and 10 mg/kg, i.p., 7 days, 1 dose/day) as measured after an acute ketamine challenge (10 mg/kg, i.p.) in adult rats in the conditioned-place preference test (CPP) **(A-C)** naïve and **(D-E)** maternally-deprived (MD) rats. Data represent mean ± SEM of the % time spent in the paired chamber **(A, D)**, the number of entries in the paired chamber **(B, E)**, and the distance (cm) traveled during the test **(C, F)**. Individual values are shown for each rat. Two-way ANOVAs (independent variables: Sex and Treatment) are shown in Supplementary Table S3. Significant effects of Sex: #*p* < 0.05 when comparing male vs. female rats. \*\**p* < 0.01 and \**p* < 0.05 vs. the corresponding Vehicle (V) group.

Interestingly, when ketamine was used to challenge rats previously exposed to maternal separation early in life and treated with ketamine in adolescence, significant Sex x Treatment interactions were detected both for the % time spent in the paired chamber (Fig. 6D), and the number of entries (Fig. 6E; see Supplementary Table S3). *Post-hoc* comparisons revealed that while a prior adolescent ketamine exposure decreased the conditioned-response induced by adult ketamine in female rats (1 mg/kg: -12 ± 4% less time in paired chamber; \**p* = 0.024; 10 mg/kg: -11 ± 4% time in paired chamber; \**p* = 0.033; and -8 ± 3% entries in paired chamber; \**p* = 0.036; Fig. 6D-E), it increased the one observed in male rats (5 mg/kg: +9 ± 3 entries in paired chamber; \**p* = 0.016; 10 mg/kg: +10 ± 3% entries in paired chamber; \*\**p* = 0.005; Fig. 6E) when compared to the corresponding vehicle-treated group.

## Discussion

Overall, the present results reinforce the potential for ketamine to induce signs of antidepressant-like efficacy in adolescent rats for both sexes but at different dose ranges. However, some safety concerns seemed associated with these beneficial effects; although adolescent ketamine did not induce reinforcing-like features in adolescence, it stimulated psychomotor-like responses with signs of sensitization following a repeated treatment in adolescent female rats. Interestingly, adolescent ketamine did not affect long-term cognitive performance in adulthood. However, it changed the reinforcing-like properties induced by a challenge dose of ketamine in adulthood, but in a different way for each sex, while it increased the response in male rats, a decreased response was observed for females.

The antidepressant-like effects of ketamine were tested under stressful test-conditions in naïve and maternally-deprived rats of both sexes. While efficacious effects were observed for both sexes, but at different dose-ranges (acute vs. the need for a repeated paradigm), in naïve rats, the present results showed a lack of response in maternally-deprived rats. In particular, while a higher dose of ketamine was enough to induce an acute response in female rats, for the lower doses tested, repetitive administrations were needed to show efficacy in adolescent naïve rats of both sexes. As previously reported, rats exposed to early-life stress (i.e., maternal deprivation) might require higher and/or longer treatment paradigms to induce a beneficial response (see [51,35]). In particular, although the model of early-life stressed used in the present study is moderate in terms of inducing basal changes in affective-like behavior in adolescence, it has proven useful for evaluating the influence of stress at early ages on pharmacological responses without the need to induce a pro-depressant-like phenotype [52,35]. Moreover, this maternal separation paradigm induced signs of hippocampal neurotoxicity in adolescent rats [34], as well as long-term decays in cognitive performance, as characterized in the present study. In fact, our novel results showed that adult rats exposed to maternal separation spent more time finding the escape box as compared to naïve rats in the Barnes maze test. The data suggested a stronger impact on male rats, consistently with studies showing that male rats were more vulnerable to this type of early life-stress when compared to females (e.g., [53]). Therefore, higher doses of ketamine might be needed to observe an antidepressant-like response in adolescent rats exposed to early-life stress. Up to here, and in line with the prior studies (e.g., [41,51,35]), our results prove that adolescent ketamine induced signs of efficacy for both sexes but at different dose-ranges and in different behavioral tests. Studies assessing sex differences in the antidepressant-like response induced by ketamine in adolescence are scarce, but the present results aligned with prior studies in adult rats showing that female rodents are more sensitive to ketamine-induced effects in the context of affective-like responses (e.g., [54-55]). The observed sex differences induced by ketamine might be related to the role of sex hormones, since several studies have shown that estrogens play an important role in the antidepressant-like response to ketamine in adult rats (e.g., [54,36]).

One of the main concerns about using ketamine during adolescence is related to its short- and long-term safety prolife, which has not been fully characterized yet. One of the primary issues is regarding ketamine’s abuse liability (e.g., [18-19,56]). In this context, in the present study, we assessed the reinforcing properties of ketamine during adolescence, at the same age-window in which we evaluated its antidepressant-like potential, and exclusively in naïve rats (doses of 5 and 10 mg/kg, acute and repeated effects) in line with the observed antidepressant-like responses. Our results suggested that the doses of ketamine used to induce an antidepressant-like effect in adolescence did not induce reinforcing-like responses in the conditioned place preference, at least with the short conditioning-paradigm tested [47]. Therefore, one could not exclude those other conditioning paradigms, based on longer conditioned phases, might induce a conditioned response for ketamine, in line with the results observed in adult rodents (e.g., [24]).

Another aspect evaluated during adolescence was the psychomotor-like effects induced by the highest dose tested of ketamine (10 mg/kg, acute and repeated effects) in naïve rats, simulating the administration paradigm of the observed antidepressant-like response. The results showed that ketamine, both acutely and after a repeated treatment of 7 doses, increased the overall distance travelled by rats. In particular, the effects were more pronounced in female rats and were time-dependent, since the distance traveled normalized 15 minutes post-injection. Given that rats were scored in the forced-swim test 30 min post-injection (once locomotion was back to normal), and that antidepressant-like responses were also observed in the novelty-suppressed feeding test (while no changes were present in distance travelled), we reinforced that the antidepressant-like effects of ketamine were not caused by its increase in locomotor activity. Moreover, ketamine-induced hyperlocomotion was sex-specific, as it observed in female rats, in line with prior studies showing that females seemed more sensitive to ketamine-induced effects on locomotion [57-59]. The greater sensitivity of females to the locomotor effects of ketamine could be due to sex differences observed in the metabolism of ketamine and other phencyclidines [57]. Furthermore, after the repeated treatment with ketamine (7 doses of ketamine, 1 dose daily), female rats showed signs of psychomotor sensitization, a characteristic of substances with addictive-like potential (e.g., [60]). Prior reports also described sex differences in the locomotor sensitization response induced by ketamine [61-62], denotating a potential addictive-like liability in adolescence that deserves some caution and/or further studies before using ketamine as an antidepressant at this early age.

Finally, we evaluated the long-term safety profile in adulthood following adolescent ketamine treatment, both at the level of cognitive performance and addiction-liability (i.e., rewarding response to a ketamine challenge dose). Adolescent ketamine, in line with other clinical data [63,16], showed a good safety prolife when administered at subanesthetic doses in terms of not altering cognition in the Barnes maze test. Similarly, another study also demonstrated no changes in cognitive performance in the Barnes maze following a repeated treatment with ketamine (at higher doses than the ones used here) during adolescence [64]. Other experiments regarding the impact of ketamine on cognition came inconsistent probably due to differences in treatment duration, dose used, age of treatment initiation, etc. (e.g., see more details as reviewed by [65]).

On a negative note, adolescent ketamine induced changes in the reinforcing properties of ketamine in adulthood as evaluated in the conditioned place preference when rats were re-exposed to a challenge dose of ketamine (10 mg/kg). The response was sex-dependent and only present when rats were also previously exposed to an early-life stressor (i.e., maternal deprivation). In particular, the rewarding-like potential of ketamine was exacerbated in adult male rats with a history of adolescent ketamine, while it was diminished in females. These effects were not observed in naïve rats, therefore suggesting, in line with the multiple-hit hypothesis, that the accumulation of vulnerability factors (i.e., maternal separation, adolescent ketamine, male vulnerability, and drug re-exposure in adulthood) might be behind the negative effects observed in male rats. Prior several studies already showed sex differences in the vulnerability to abuse liability by subanesthetic treatments with ketamine (reviewed by [66]). Similar to the increased male vulnerability, another study in mice showed that a repeated treatment with ketamine during adolescence modified the reinforcing properties of other drugs of abuse, such as cocaine, in adulthood, and did so exclusively in males [67]. The results demonstrated that the long-term impact of adolescent ketamine was sex-specific, being more deleterious in male rats. Therefore, the present addictive-like vulnerability observed for adult male rats together with the psychomotor-like sensitization induced by ketamine in adolescent female rats suggests a bit of caution and the need for further research before moving forward the use of ketamine as an antidepressant for adolescence.

## Role of the Funding Source

Funding for this study was provided by PID2020-118582RB-I00 and PID2023-151640OB-I00 (MCIN/AEI/10.13039/501100011033) and partially sponsored and promoted by the Comunitat Autònoma de les Illes Balears through the Servei de Recerca i Desenvolupament and the Conselleria d’Educació i Universitats (PDR2020/14 - ITS2017-006) to MJG-F. The program JUNIOR (IdISBa, GOIB) supported SL-C’s salary. JJ-P. was funded by a predoctoral scholarship from Conselleria de Fons Europeus, Universitat i Cultura, Govern de les Illes Balears (FPI_022_2022).

## Contributors

**Jordi Jornet-Plaza** - Conceptualization; Data Curation; Formal Analysis; Methodology; Original Draft Preparation; Writing - Review & Editing.

**Sandra Ledesma-Corvi** - Formal Analysis; Methodology; Writing - Review & Editing.

**M. Julia García-Fuster** - Conceptualization; Data Curation; Funding Acquisition; Project Administration; Resources; Original Draft Preparation; Writing - Review & Editing.

All authors contributed to and have approved the final manuscript.

## Conflict of Interest

All authors declare that they have no conflicts of interest.

## Data availability

Raw data will be made available upon request to the corresponding author.

## Supplementary Materials

Supplementary material associated with this article can be found, in the online version.

## Notes

### Competing Interest Statement

The authors have declared no competing interest.

