## Supplemental Material for "Safety concerns following the use of ketamine as a potential antidepressant for adolescent rats of both sexes"

### Supplementary Materials

**Table. S1.** Two-way ANOVAs analyses (intendent variables: Sex and Treatment), both including F (DFn, DFd) and *p* values, for data represented in Fig. 2-3. Green-shadow boxes represent statistically significant comparisons.

**Affective-like responses in adolescece**

| | Acute effects D1 - FST | Sex | Treatment | Sex x Treatment |
|---|---|---|---|---|
| Naïve rats | Immobility (s) | F (1, 125) = 4.824; #*p* = 0.023 | F (3, 125) = 5.413; *p* = 0.002 | F (3, 125) = 5.035; ; p = 0.003 |
| | Climbing (s) | F (1, 125) = 4.264; #p = 0.041 | F (3, 125) = 4.921; p = 0.003 | F (3, 125) = 5.008; p = 0.003 |
| | Swimming (s) | F (1, 125) = 0.197; p = 0.658 | F (3, 125) = 3.511; p = 0.017 | F (3, 125) = 1.681; p = 0.175 |
| MD rats | Immobility (s) | F (1, 120) = 0.014; p = 0.905 | F (3, 120) = 0.204; p = 0.893 | F (3, 120) = 2.785; p = 0.044 |
| | Climbing (s) | F (1, 120) = 0.964; p = 0.328 | F (3, 120) = 1.106; p = 0.350 | F (3, 120) = 3.781; p = 0.012 |
| | Swimming (s) | F (1, 120) = 5.141; #p = 0.025 | F (3, 120) = 2.892; #p = 0.038 | F (3, 120) = 2.418; p = 0.070 |
| | **Repeated effects D8 - FST** | **Sex** | **Treatment** | **Sex x Treatment** |
| Naïve rats | Immobility (s) | F (1, 123) = 25.69; *### p<0.001* | F (3, 123) = 1.772; p = 0.156 | F (3, 123) = 1.379; p = 0.2524 |
| | Climbing (s) | F (1, 123) = 25.63; ### p<0.001 | F (3, 123) = 3.266; p =0.024 | F (3, 123) = 1.153; p = 0.330 |
| | Swimming (s) | F (1, 123) = 3.946; #p = 0.049 | F (3, 123) = 2.974; p = 0.034 | F (3, 123) = 1.042; p = 0.377 |
| MD rats | Immobility (s) | F (1, 121) = 3.211; p=0.076 | F (3, 121) = 0.089; P=0.966 | F (3, 121) = 1.613; p = 0.190 |
| | Climbing (s) | F (1, 121) = 2.203; p = 0.140 | F (3, 121) = 0.633; P=0.595 | F (3, 121) = 0.998; p = 0.396 |
| | Swimming (s) | F (1, 121) = 3.245; p = 0.074 | F (3, 121) = 1.448; p = 0.232 | F (3, 121) = 2.738; p = 0.046 |
| | **Repeated effects D12 - NSFT** | **Sex** | **Treatment** | **Sex x Treatment** |
| Naïve rats | Feeding time (s) | F (1, 125) = 0.487; p = 0.487 | F (3, 125) = 5.998; p <0.001 | F (3, 125) = 1.235; p = 0.300 |
| | Latency to center (s) | F (1, 125) = 1.334; p = 0.250 | F (3, 125) = 2.210; p = 0.090 | F (3, 125) = 0.401; p = 0.753 |
| MD rats | Feeding time (s) | F (1, 123) = 1.950; p = 0.165 | F (3, 123) = 5.765; p = 0.001 | F (3, 123) = 1.487; p = 0.221 |
| | Latency to center (s) | F (1, 123) = 0.951; p = 0.331 | F (3, 123) = 2.047; p = 0.111 | F (3, 123) = 0.648; p = 0.586 |

**Conditioned-place preference in adolescence**

| | Acute effects D1 - CPP | Sex | Treatment | Sex x Treatment |
|---|---|---|---|---|
| Naïve rats | Paired chamber (% time) | F (1, 34) = 0.309; p = 0.582 | F (2, 34) = 3.103; p = 0.056 | F (2, 34) = 0.067; p = 0.936 |
| | Paired chamber (entries) | F (1, 34) = 2.290; p = 0.139 | F (2, 34) = 1.940; p = 0.159 | F (2, 34) = 0.863; p = 0.431 |
| | Distance (cm) | F (1, 34) = 0.262; p = 0.612 | F (2, 34) = 0.531; p = 0.593 | F (2, 34) = 0.307; p = 0.737 |
| | **Repeated effects D8 - CPP** | **Sex** | **Treatment** | **Sex x Treatment** |
| Naïve rats | Paired chamber (% time) | F (1, 34) = 1.159; p = 0.289 | F (2, 34) = 0.608; p = 0.550 | F (2, 34) = 0.391; p = 0.679 |
| | Paired chamber (entries) | F (1, 34) = 5.731; #p = 0.022 | F (2, 34) = 1.123; p = 0.337 | F (2, 34) = 0.040; p = 0.960 |
| | Distance (cm) | F (1, 34) = 10.520; ##p = 0.003 | F (2, 34) = 4.081; p = 0.026 | F (2, 34) = 0.577; p = 0.567 |

**Table. S2.** Three-way ANOVAs analyses (intendent variables: Sex, Treatment and Time or Day), both including F (DFn, DFd) and *p* values, for data represented in Fig. 4. Green-shadow boxes represent statistically significant comparisons.

**Psychomotor sensitization in adolescence**

| | Acute effects D1 - Open Field | Sex | Treatment | Sex x Treatment | Time | Time x Treatment | Sex x Treatment x Time |
|---|---|---|---|---|---|---|---|
| Naïve rats | Distance (cm) | F (1, 276) = 6.065; #p= 0.014 | F (1, 276) = 51.230; p <0.001 | F (1, 276) = 8.349; p = 0.004 | F (11, 276) = 4.088; p <0.001 | F (11, 276) = 1.420; p = 0.163 | F (11, 276) = 1.345; p = 0.199 |
| | **Repeated effects D7 - Open Field** | **Sex** | **Treatment** | **Sex x Treatment** | **Time** | **Time x Treatment** | **Sex x Treatment x Time** |
| Naïve rats | Distance (cm) | F (1, 276) = 50.550; ###p < 0.001 | F (1, 276) = 18.720; p < 0.001 | F (1, 276) = 14.650; p < 0.001 | F (11, 276) = 6.840; p < 0.001 | F (11, 276) = 2.652; p = 0.003 | F (11, 276) = 2.029; p = 0.026 |
| | **D1 vs. D7 - Sensitization** | **Sex** | **Treatment** | **Sex x Treatment** | **Day** | **Day x Treatment** | **Sex x Treatment** |
| Naïve rats | Total distance (cm) | F (1, 23) = 8.253; p = 0.009 | F (1, 23) = 30.17; p < 0.001 | F (1, 23) = 7.385; p = 0.012 | F (1, 23) = 8.937; p = 0.007 | F (1, 23) = 3.764; p = 0.065 | F (1, 23) = 7.385; p = 0.012 |

**Fig. S1. Long-term effects of maternal separation (PND 9-10) in the performance observed in the Barnes maze in adulthood.** Time (s) utilized to resolve the Barnes maze during training sessions (A), Test 1 (B) and Test 2 (novel cue conditions) (C). While each training session lasted 180 s, test sessions lasted 90 s (see more details in the Experimental procedures section of the manuscript). A three-way ANOVA detected a significant effect of Early-life Condition (Training session: F (1, 30) = 8.14; *p* = 0.008), demonstrating an overall longer time (s) needed for rats exposed to maternal separation (MD), and independently of sex, to resolve the maze during training. Moreover, two-way ANOVAs also detected significant effects of Early-life Condition for Test 1 (F (1, 30) = 7.64; *p* = 0.010) and Test 2 (F (1, 30) = 12.46; *p* = 0.001), as well as significant interactions between Sex x Early-life Condition (Test 1: F (1, 30) = 7.75; *p* = 0.009 and Test 2: F (1, 30) = 5.26; *p* = 0.029). Šídák's multiple comparisons tests revealed that early-life stress induced a more profound impact in male rats, which showed significant longer times to resolve the maze as compared to naïve rats of the same sex (Test 1: +42 ± 11 s, ****p* < 0.001; Test 2: +42 ± 10 s, ****p* < 0.001).

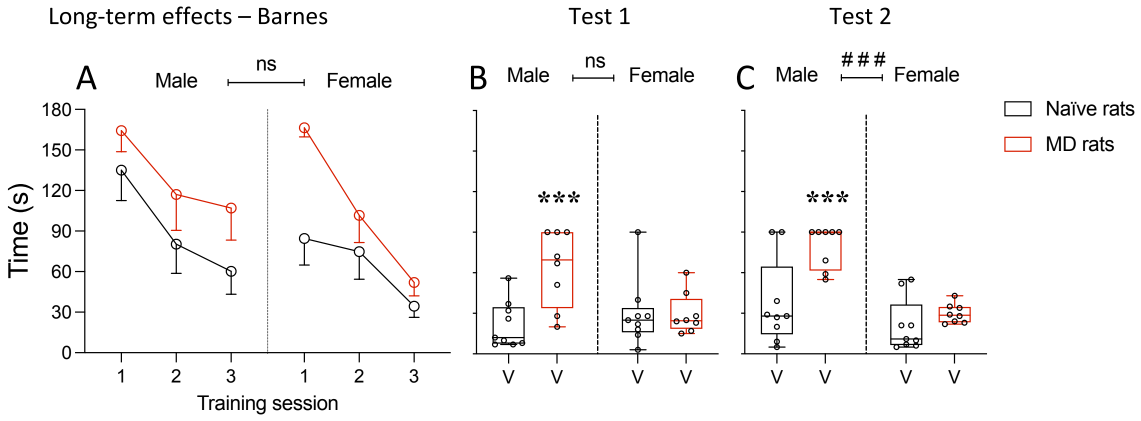


**Table. S3.** Two-way ANOVAs analyses (intendent variables: Sex and Treatment), both including F (DFn, DFd) and *p* values, for data represented in Fig. 5-6. Green-shadow boxes represent statistically significant comparisons.

**Barnes maze in adulthood**

| | Time (s) | Session | Treatment | Session x Treatment |
|---|---|---|---|---|
| **Naïve rats** | Male rats | F (1.955, 56.70) = 34.28; p < 0.001 | F (3, 29) = 0.567; p = 0.641 | F (6, 58) = 0.731; p = 0.626 |
| | Female rats | F (1.790, 53.71) = 6.237; p = 0.005 | F (3, 30) = 0.225; p = 0.878 | F (6, 60) = 0.512; p = 0.797 |
| **MD rats** | Male rats | F (1.723, 49.97) = 19.94; p < 0.001 | F (3, 29) = 0.763; p = 0.524 | F (6, 58) = 0.576; p = 0.748 |
| | Female rats | F (1.750, 50.76) = 16.30; p < 0.001 | F (3, 29) = 2.855; p = 0.054 | F (6, 58) = 1.196; p = 0.322 |
| | **Test performance** | **Sex** | **Treatment** | **Sex x Treatment** |
| **Naïve rats** | Test 1 | F (1, 59) = 1.956; p = 0.167 | F (3, 59) = 1.136; p = 0.342 | F (3, 59) = 0.886; p = 0.458 |
| | Test 2 | F (1, 59) = 5.469; p = 0.029 | F (3, 59) = 0.638; p = 0.594 | F (3, 59) = 0.646; p = 0.589 |
| **MD rats** | Test 1 | F (1, 58) = 1.961; p = 0.167 | F (3, 58) = 2.060; p = 0.115 | F (3, 58) = 1.641; p = 0.190 |
| | Test 2 | F (1, 58) = 3.768; p = 0.057 | F (3, 58) = 0.241; p = 0.869 | F (3, 58) = 3.346; p = 0.025 |

**Conditioned-place preference in adulthood**

| | CPP | Sex | Treatment | Sex x Treatment |
|---|---|---|---|---|
| **Naïve rats** | Paired chamber (% time) | F (1, 55) = 0.026; p = 0.873 | F (3, 55) = 1.672; p = 0.184 | F (3, 55) = 1.491; p = 0.227 |
| | Paired chamber (entries) | F (1, 55) = 0.051; p = 0.822 | F (3, 55) = 0.933; p = 0.431 | F (3, 55) = 2.120; p = 0.108 |
| | Distance (cm) | F (1, 55) = 6.788; p = 0.012 | F (3, 55) = 1.998; p = 0.125 | F (3, 55) = 4.875; p = 0.005 |
| | **CPP** | **Sex** | **Treatment** | **Sex x Treatment** |
| **MD rats** | Paired chamber (% time) | F (1, 57) = 0.006; p = 0.938 | F (3, 57) = 0.908; p = 0.443 | F (3, 57) = 2.935; p = 0.041 |
| | Paired chamber (entries) | F (1, 57) = 0.817; p = 0.370 | F (3, 57) = 0.195; p = 0.899 | F (3, 57) = 6.351; p < 0.001 |
| | Distance (cm) | F (1, 57) = 6.329; p = 0.015 | F (3, 57) = 0.194; p = 0.900 | F (3, 57) = 0.953; p = 0.421 |
